# SegVeg: Segmenting RGB images into green and senescent vegetation by combining deep and shallow methods

**DOI:** 10.1101/2022.03.24.485604

**Authors:** Mario Serouart, Simon Madec, Etienne David, Kaaviya Velumani, Raul Lopez Lozano, Marie Weiss, Frédéric Baret

## Abstract

The pixels segmentation of high resolution RGB images into background, green vegetation and senescent vegetation classes is a first step often required before estimating key traits of interest including the vegetation fraction, the green area index, or to characterize the sanitary state of the crop. We developed the SegVeg model for semantic segmentation of RGB images into the three classes of interest. It is based on a U-net model that separates the vegetation from the background. It was trained over a very large and diverse dataset. The vegetation pixels are then classified using a SVM shallow machine learning technique trained over pixels extracted from grids applied to images. The performances of the SegVeg model are then compared to a three classes U-net model trained using weak supervision over RGB images with predicted pixels by SegVeg as groundtruth masks.

Results show that the SegVeg model allows to segment accurately the three classes, with however some confusion mainly between the background and the senescent vegetation, particularly over the dark and bright parts of the images. The U-net model achieves similar performances, with some slight degradation observed for the green vegetation: the SVM pixel-based approach provides more precise delineation of the green and senescent patches as compared to the convolutional nature of U-net. The use of the components of several color spaces allows to better classify the vegetation pixels into green and senescent ones. Finally, the models are used to predict the fraction of the three classes over the grids pixels or the whole images. Results show that the green fraction is very well estimated (R^2^=0.94) by the SegVeg model, while the senescent and background fractions show slightly degraded performances (R^2^=0.70 and 0.73, respectively).

We made SegVeg publicly available as a ready-to-use script, as well as the entire dataset, rendering segmentation accessible to a broad audience by requiring neither manual annotation nor knowledge, or at least, offering a pre-trained model to more specific use.

## 1 Introduction

The fraction of vegetation (VF) is a key trait that drives the partitioning of radiation between the background and the vegetation. It is used in several studies as a proxy of crop state [1] and yield [2, 3]. The complement to unity of VF is the gap fraction that is used to estimate the plant area index. However, several eco-physiological processes such as photosynthesis and transpiration are driven by the amount of green surfaces that exchange mass and energy with the atmosphere. GF is also used to estimate the green area index (GAI) [4] defined as the area of green vegetation elements per unit horizontal ground area. GF is a more relevant trait that should be used when describing crop functioning [5]. The difference between VF and GF is the senescent fraction (SF=VF-GF), sometimes called non-photosynthetic fraction [6]. For crops, SF depends on both the growth stage and state of the plants. The SF trait is used to characterize a biotic or abiotic stress, to describe nutrient recycling, and monitor the ageing process [7–9]. Some studies focused on the ability of genotypes to stay green by delaying senescence and potentially improve productivity [10, 11].

Several remote sensing methods have been developed to estimate GF and SF using the spectral variation of the signal observed at the canopy scale from metric to decametric pixels [12]. VF, GF and SF can be also computed using very high spatial resolution images with pixels from a fraction of mm to a cm, i.e., significantly smaller than the typical dimension of the objects (plants, organs). Identifying the green and senescent parts of the plants provides additional information for organ and disease detection or characterization. RGB cameras with few millions to tens of millions of pixels are currently widely used as non-invasive high throughput techniques applied to plant breeding, farm management, and yield prediction [13–15]. These cameras are borne on multiple platforms, including drones [16], ground vehicles [17], hand-held systems [18] or set on a fixed pod [15].

Several methods have been proposed to identify the green pixels into RGB images including thresholding color indices [19], and machine learning classification [20] based on few color-space representations. However, these techniques are limited at least by two main factors:

### Confounding effects

Depending on the illumination conditions and on the quality of the optics of the camera, part of the soil may appear green due to chromatic aberration. Further, parts of the image that are saturated, with strong specular reflection or very dark will be very difficult to classify using only the color of the pixel. Finally, the soil may also show greenish when it contains algae [21].

### Continuity of colors

At the cellular scale, senescence results from the degradation of pigments that generally precedes cell death [22]. During the degradation process, changes in the pigment composition results into a wide palette of leaf color in RGB imagery, with a continuity between “green” and “senescent” states. Further, when pixels are located at the border of an organ, its color will be intermediate between that of the organ and that of the background. This problem is obviously enhanced when the spatial resolution of the RGB image is too coarse.

It is therefore difficult to segment accurately and robustly the green vegetation parts of a RGB image using only the color information of the pixels. The same limitations apply to the segmentation of the vegetation senescent parts. In addition, crop residues on the background are difficult to distinguish from the senescent vegetation observed on standing up plants: they show very similar range of brownish colors. Textural and contextual information should therefore be exploited to better segment RGB images into green and senescent vegetation parts.

Semantic segmentation [23] that assigns a class to each pixel of the image appears to be an attractive approach. It is based on deep learning techniques and has been applied to several domains including urban scene description for autonomous vehicles, medical imagery [24], and agriculture [25, 26]. However, images need to be labelled exhaustively into the several classes, which requires large annotation resources [27].

The objective of this paper is to develop and evaluate a two-step semantic segmentation approach called SegVeg. It labels each pixel of a very high resolution RGB image of vegetation scenes into three classes: Background, Green vegetation and Senescent vegetation. It is designed to reduce the annotation effort by combing a convolutional neural network (CNN) that splits the image into vegetation (including both green and senescent pixels) and background, to a simple support vector machine (SVM) technique that classifies the vegetation pixels into green and senescent ones. SegVeg will be compared to a CNN classifier that will directly identifies background, green vegetation, and senescent vegetation pixels. The training and testing of the models is applied to a comprehensive annotated dataset of RGB images taken from ground level over a wide range of crops and conditions.

## 2 Materials and Methods

### 2.1 The SegVeg model

The SegVeg model is made of two stages (Fig 1). In the first stage, the whole image is classified into Vegetation/Background mask using a U-net type Deep Learning network [28]. Then, the vegetation pixels are classified into Green/Senescent vegetation using a SVM.

**Figure 1:**
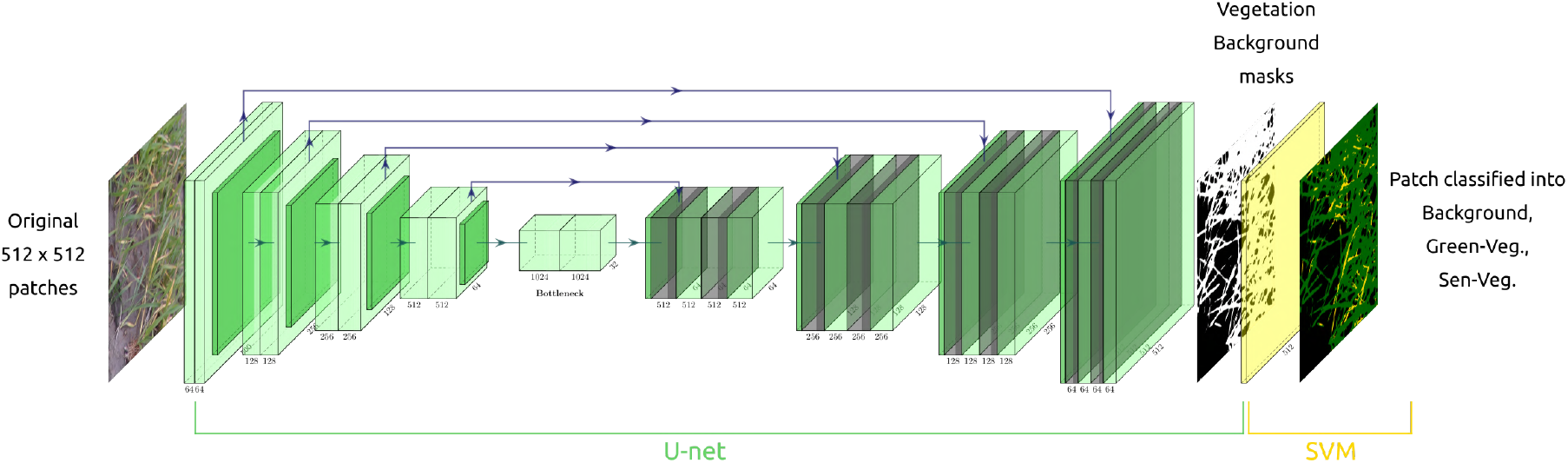
Illustration of the SegVeg architecture. The first stage is a U-net model that predicts vegetation and background masks. The second stage is a SVM that classifies the vegetation mask into green and senescent pixels. The two stages were trained over two independent datasets.

#### 2.1.1 First stage: vegetation-background segmentation

U-net is a deep learning model with encoder-decoder network architecture that is widely used for image semantic segmentation. The model was trained to predict two classes: Vegetation (green or senescent) and Background. Efficientnet-B2 architecture [29] with weights initialized on ImageNet was used as the backbone architecture. Patches of 512 × 512 pixels were used for training after data augmentation based on the Albumentations library [30]. The training database used for this first stage is described in a next section. The training process was based on a Dice loss function with an Adam optimizer and a progressive learning rate using 100 epochs split in 3 batches. The Python Segmentation library under PyTorch was used [31] with GPU activation (GeForce RTX 3090).

#### 2.1.2 Second stage: classification of green and senescent vegetation pixels

The Support Vector Machine (SVM) is an efficient machine learning classification method widely used for image segmentation [32–34]. It maps the original features space to some higher-dimensional space where the training dataset is separable. Several color spaces and transformations [35] were used to classify pixels including (RGB, HSV, CIELab, Grayscale, Luminances, CMYK, YCbCr, YIQ) derived from the original RGB values. A total of 23 potential input features were thus computed, namely R, G, B; H, S, V; L, a, b; GE; LA, LB, LC; C, M, Y, K; Yi, Cb, Cr; Yj, I, and Q. However, the possible redundancy and irrelevancy of some features may decrease the accuracy of the classification. We then selected the most appropriate inputs using the step forward wrapper method [36]. Finally, 14 input features were kept: R, G, B, H, S, a, b, GE, M, YE, Cb, Cr, I, and Q. This SVM second stage was calibrated over a specific pixels grid dataset described later. The hyperparameters were tuned using a grid search algorithm leading to the optimal values: C: 1, *γ*: 10^*−*3^, kernel: rbf.

### 2.2 The 3-classes U-net model

A three classes U-net model was used as a reference to evaluate the proposed SegVeg model. However, since we did not have a dataset of images annotated into the three classes (background, green vegetation, senescent vegetation), we applied the SegVeg model to the dataset used to train U-net in the first stage of the SegVeg model. The U-net architecture and hyper-parameters used for the first stage of the SegVeg model were also employed here.

### 2.3 Training and testing datasets

#### 2.3.1 Dataset used to train the first stage of the SegVeg

Height experiments were compiled to get a wide range of acquisition conditions, species, crop states, and stages (Table 1).

**Table 1:** Characteristics of the sub-datasets used to acquire images composing the final dataset.

| Sub-datasets | Country | Year | Crops | Stage | Reference |
| --- | --- | --- | --- | --- | --- |
| **Utokyo** | Japan | 2019<br>2012 | Rice, Wheat | Vegetative | [37, 38] |
| **P2S2** | France<br>Belgium | 2018 | Wheat, Rapeseed, Sugarbeet, Potato<br>Maize, Grassland, Sunflower, Rice, Soya | All | [39] |
| **Wuhan** | China | 2012<br>2015 | Cotton, Maize, Rice | Vegetative | [40] |
| **CVPPP 1 & 2** | Italy | 2012<br>2013 | Arabidopsis, Tobacco | All | [41, 42] |
| **GEVES** | France | 2020 | Maize | Vegetative | - |
| **Phenofix** | France | 2020 | Maize | All | - |
| **Phenomobile** | France | 2020 | Wheat | Early | - |
| **Bonirob** | Germany | 2016 | Sugarbeet | Early | [43] |

The images were acquired with several cameras equipped with different focal length optics and at variable distances from the ground. All blurred images or those with poor quality were excluded from our study. The original images were then split into several 512× 512 pixels square patches, a size selected to keep enough features. A total of 2015 patches were extracted, showing a large diversity as illustrated in Table 2. The ground sampling distance (GSD) was always in between 0.3 to 2 mm to capture enough details (Fig 2).

**Table 2:** Characteristics of the sub-datasets used to compose the training dataset. UGV means Unmanned Ground Vehicle.

| Sub-datasets | Platform | Camera | Image Size (px) | Distance to ground (m) | GSD (mm) | Nb. Images |
| --- | --- | --- | --- | --- | --- | --- |
| **UTokyo** | Gantry | Canon EOS | 5184x3456 | 1.5-1.8 | 0.2-0.6 | 534 |
|  |  | Kiss X5 Garden | 1280x1024 |  |  |  |
| **P2S2** | Handheld | SONY ILCE-5000 | 5456x3632 | 2 | 0.5 | 170 |
|  |  | SONY ILCE-6000 | 6000x5000 |  |  |  |
|  |  | Canon EOS 400D | 3888x2592 |  |  |  |
|  |  | Canon EOS 60D | 5184x3456 |  |  |  |
|  |  | Canon EOS 750D | 6000x4000 |  |  |  |
| **Wuhan** | Gantry | Olympus E-450 | 3648x2736 | 0.3-5 | 0.4-0.5 | 343 |
| **CVPPP 1 & 2** | Gantry | Canon PowerShotSD1000 | 3108x2324 | 1 | 0.1-0.3 | 752 |
| **GEVES** | Handheld | SAMSUNG SM-A705FN | 3264x1836 | 2 | 0.2 | 50 |
| **Phenofix** | Gantry | SONY RX0 II | 4800x3200 | 2 | 0.6 | 30 |
| **Phenomobile** | UGV | SONY RX0 II | 4800x3200 | 1.7 | 0.8-1.4 | 76 |
| **Bonirob** | UGV | JAI AD-130GE | 1296x966 | 0.85 | 0.3 | 60 |
| **Total** |  |  |  |  |  | **2015** |

**Figure 2:**
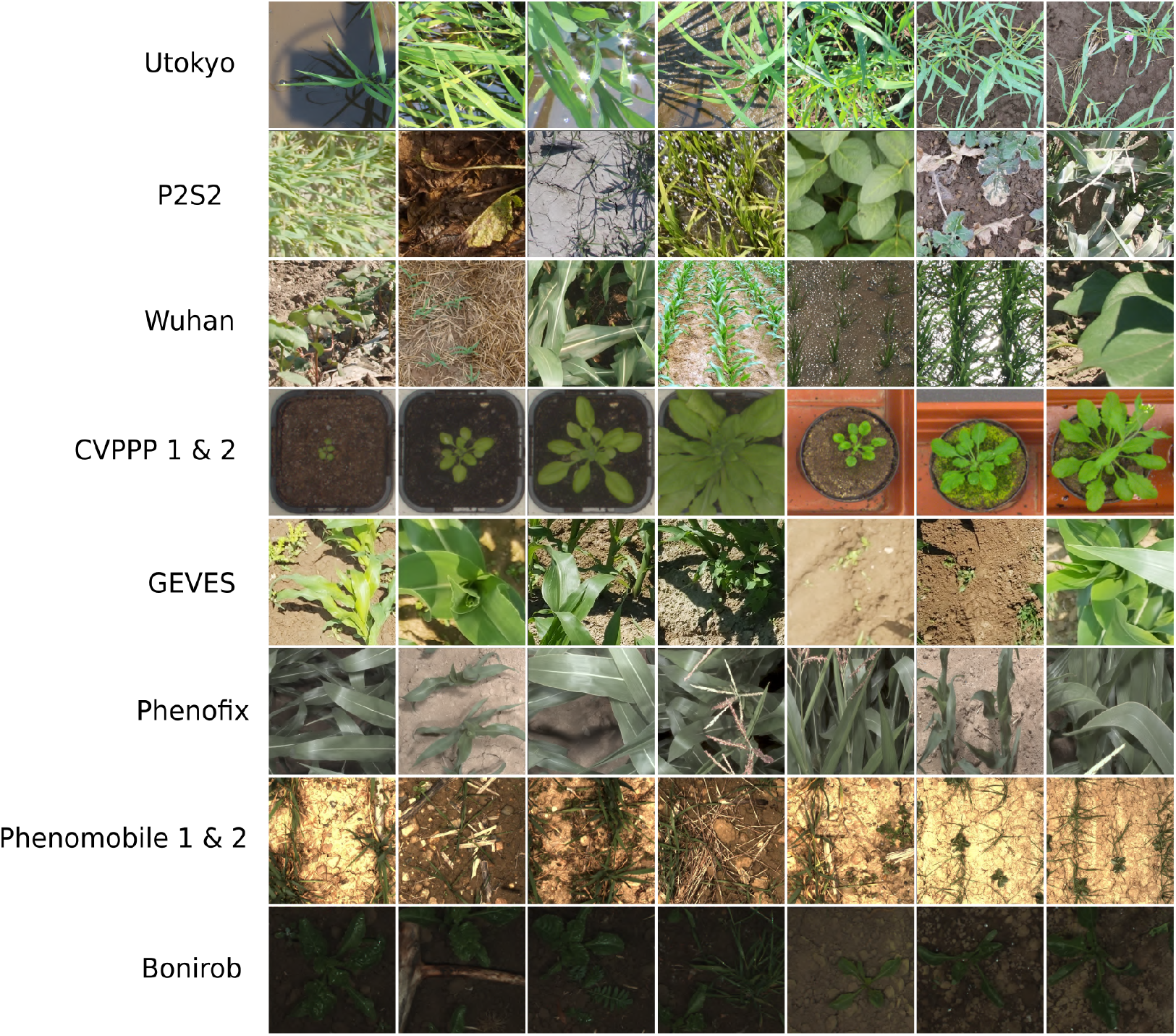
Sample of 512 × 512 pixels patches extracted from the eight experiments.

Because the image annotation is time consuming, it was subcontracted to the imageannotation. ai private company. Each original image was carefully segmented by several operators. We then verified the resulting classified images into Vegetation (Green and non-Green) and Background pixels and re-annotated the few that appeared not correct.

#### 2.3.2 Dataset used to train the second stage SVM and to evaluate the performances

The datasets described below were used to train the second stage of the SegVeg model and the three-classes U-net model. We then evaluated the performances of both the SegVeg and the three-classes U-net models.

##### 2.3.2.1 Image acquisition and extraction

Three independent datasets (LITERAL, PHENOMOBILE and P2S2) were used to train and evaluate the methods proposed. They were made of patches of 512 × 512 pixels extracted from the original RGB images, for consistency with first stage image format.

- The **LITERAL** dataset was acquired with a handheld system called LITERAL (Fig.3). An operator maintains a boom with a Sony RX0 camera fixed at its extremity. The camera looked the ground from nadir at an approximately fixed distance (Table 3). The 68 available annotated patches covered a wide range of wheat genotypes grown at several places in France, representing different growth stages and illumination conditions.
- The **PHENOMOBILE** dataset was acquired with the Phenomobile system (Fig.3), an unmanned ground vehicle [44]. This system uses flashes for image acquisition making the measurements independent from the natural illumination conditions. Images are acquired from nadir at a fixed distance from the top of the canopy (Table 3). The 173 available annotated patches covered six crops grown in four phenotyping platforms in France (Table 3).
- The **P2S2** dataset is made of 200 patches extracted from both hemispherical and nadir images. The acquisition was designed to provide a large dataset over a wide range of crops, observed under contrasted growth conditions, throughout the crop growth cycle, covering crucial phenological stages. More details on the dataset can be found in [39].

**Table 3:** Second stage dataset description.

| Datasets | LITERAL | PHENOMOBILE | P2S2 |
| --- | --- | --- | --- |
| Latitude, Longitude | 43.7°N, 5.8°E | 43.7 N°, 6.7 E° | 43.6 N°, 4.5 E° |
|  | 49.7°N, 3.0°E | 47.4 N°, 2.3 E° | 43.4 N°, 1.2 E° |
|  | 43.5°N, 1.5° E | 43.7 N°, 5.8 E° | 48.3 N°, 2.4 E° |
|  |  | 43.4 N°, 0.4 W° | 50.6 N°, 4.7 E° |
| Year | 2017-2020 | 2018-2020 | 2018 |
| Crops | Wheat | Wheat, Sunflower, Sugar beet<br>Maize, Potato, Flax | Wheat, Sunflower, Sugar beet<br>Maize, Potato, Rapeseed<br>Grassland, Rice, Soja |
| Vector | Handheld | Phenomobile | Handheld |
| Focal length (mm) | 8 | 16 – 25 | 50 |
| Camera | Sony RX0 II | Baumer VCXG-124C | ILCE-6000 SONY<br>Canon EOS 750D |
| Image size (pixels) | 4800x3200 | 4096x3000 | 6000x4000<br>3888x2592 |
| Pixel size (µm) | 2.74 | 3.45 | 3.72 |
| Distance to ground (m) | 1.5 – 2.5 | 2 – 4.5 | 1.5 – 2 |
| GSD (mm) | 0.65 | 1.3 | 0.5 |

Several cameras were used for the acquisition of the three datasets, resulting in differences in image quality and GSD (Table 3). Note that the GSD of this dataset (Table 3) is consistent with that of the previous dataset (Table 2). A total of 441 patches of 512 × 512 pixels were finally selected to represent a wide diversity (Fig. 3).

**Figure 3:**
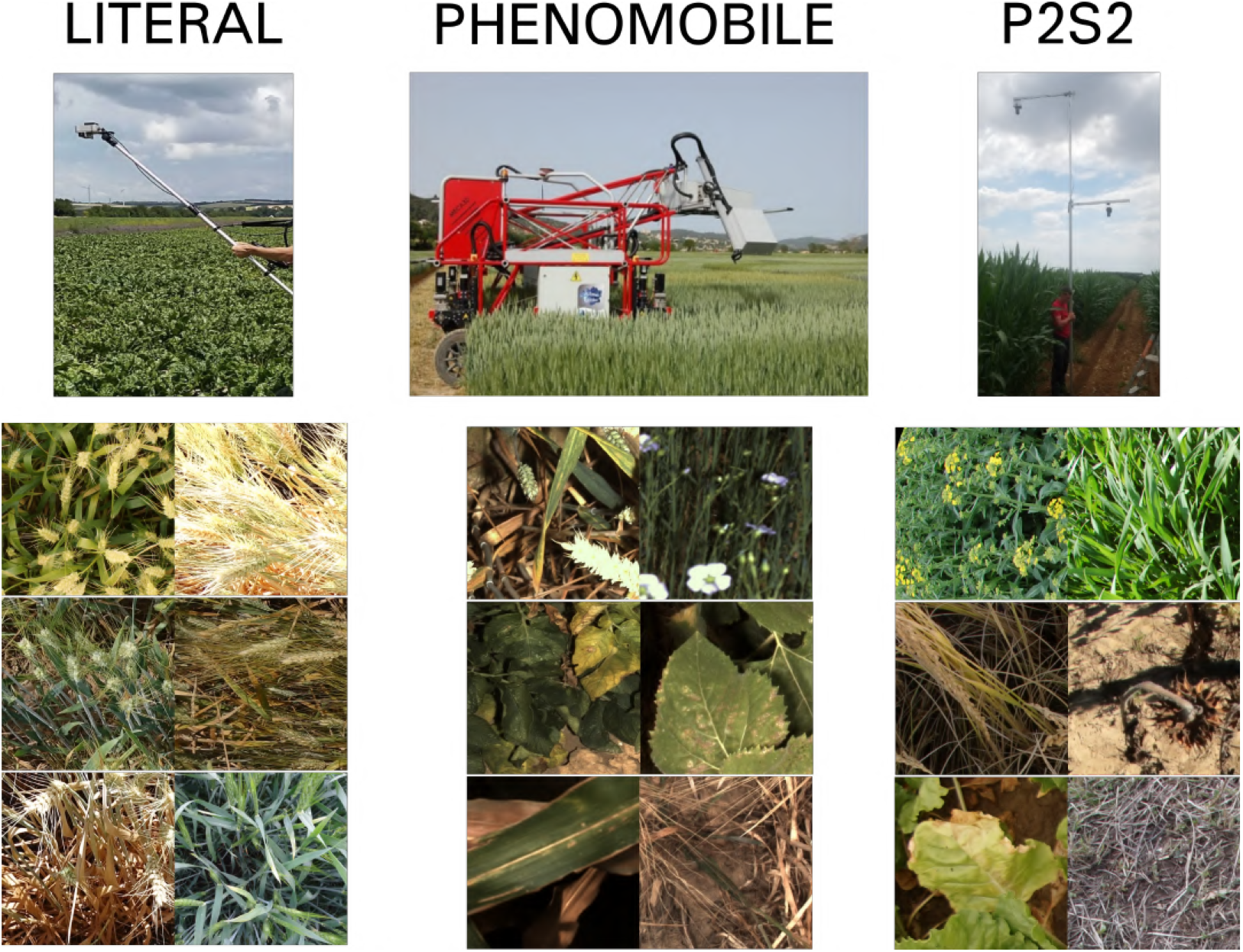
The acquisition systems used for the three datasets: LITERAL, PHENOMOBILE, and P2S2 and their respective examples of 512 × 512 images patches extracted from the three systems.

##### 2.3.2.2 Image annotation

The pixels were classified into one of the following six classes, namely: ***Green Vegetation***, ***Senescent Vegetation***, ***Background***, ***Green/Senescent Vegetation Unsure, Unknown***, ***Others***. This allowed us to remove pixels with uncertain annotations and potential bias in the training phase. The ***Green/Senescent Vegetation Unsure, Unknown*** and ***Others*** were for instance not used in the training and evaluation of the proposed models. However, because of the complexity, subjectivity and time required to assign pixels into the six classes listed above, the annotation was limited to a small number of pixels per patches. This sampled annotation is possible because the second stage of SegVeg (SVM) does not require the patches to be exhaustively annotated. We used a grid displayed on each patch, where the pixels to be classified were located at the intersection of the grid points (Fig. 4). The regular square matrix can vary from 8 to 11 pixels on a side, depending on images. The Datatorch web-based platform [45] was used to label the patches. A second round of pixel labeling was performed by several annotators over the uncertain pixels to find a better consensus on the labeling.

**Figure 4:**
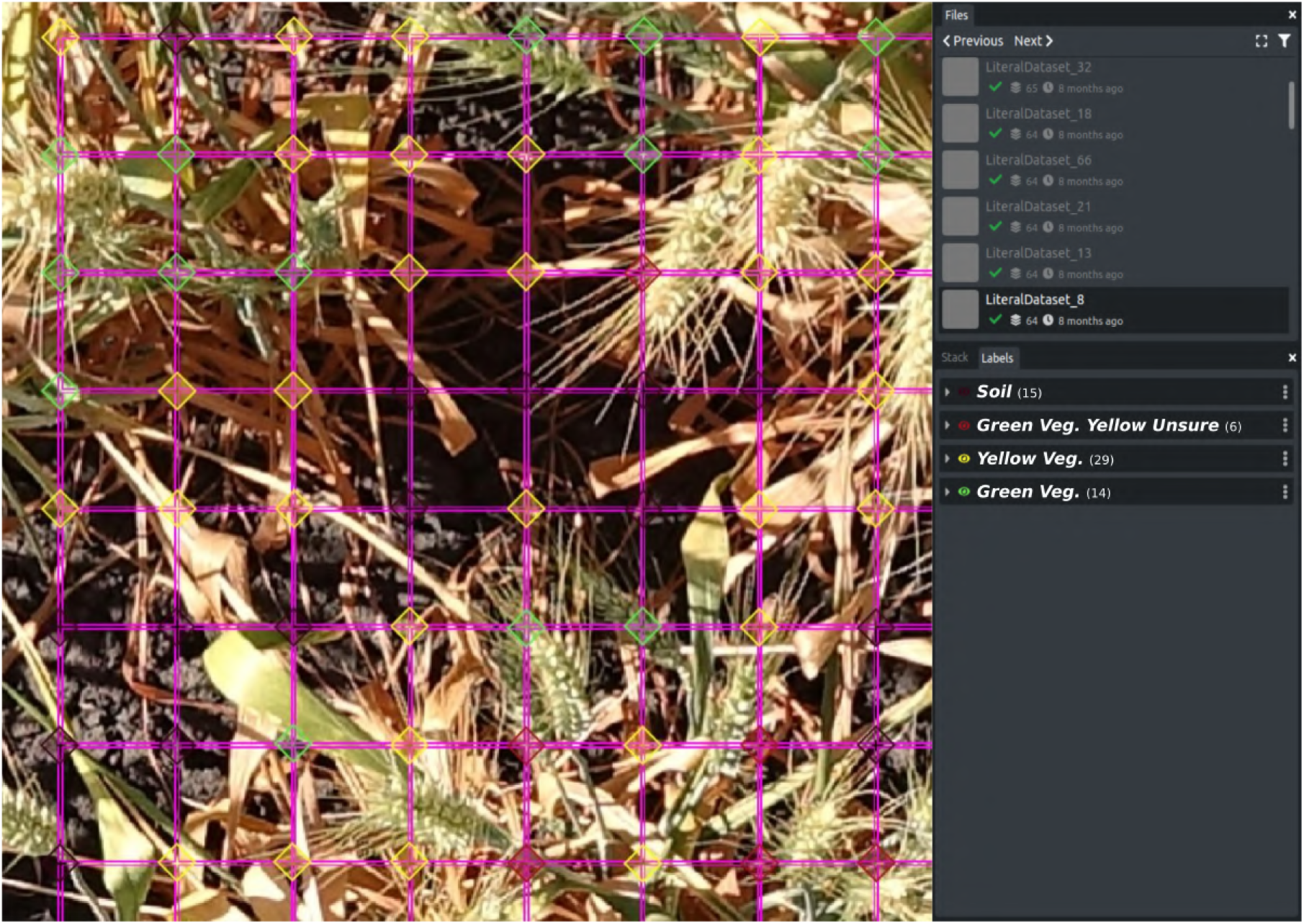
Screenshot of the used tool to annotate the pixels on the grid. The annotator classifies every pixel at the intersection of the grid to the one of the six classes: ***Green Vegetation, Senescent Vegetation***, ***Background***, ***Green / Senescent Vegetation Unsure, Unknown***, ***Others***. The diamond feature at the intersection of the grid is only used to improve the visibility. Only the single pixel located exactly at the intersection was labeled.

Among the 441 annotated grids (Table 4), the unsure classes represented about 8% of the total number of pixels, for the PHENOMOBILE dataset the integrated flashes provided better pixel interpretation leading to fewer confusions. This dataset is publicly available on Zenodo following this link https://github.com/mserouar/SegVeg (When published linked with ORCID).

**Table 4:** Distribution of labeled pixel for the three datasets.

| Datasets | Nb. of<br>labelled<br>patches | Nb. of<br>labelled<br>pixels | % Classes |  |  |  |  |  |
| --- | --- | --- | --- | --- | --- | --- | --- | --- |
|  |  |  | Green<br>Veg. | Sen.<br>Veg. | Background | Green /<br>Sen. Veg.<br>Unsure | Unknown | Other |
| **LITERAL** | 68 | 4260 | 46.5 | 15.8 | 15.0 | 13.1 | 9.5 | 0.1 |
| **PHENOMOBILE** | 173 | 8266 | 40.3 | 31.1 | 27.6 | 0.1 | 0.8 | 0.1 |
| **P2S2** | 200 | 18559 | 43.6 | 16 | 15.2 | 11.1 | 13 | 1.1 |
| **Total** | 441 | 31085 | 43.4 | 20.5 | 19.75 | 8.1 | 7.8 | 0.45 |

##### 2.3.2.3 Split between training and testing datasets

Pixels difficult to annotate, i.e. belonging to the ***Green/Senescent Vegetation Unsure, Unknown***, ***Others*** classes (Table 4), were eliminated from the training and testing datasets. A total of 19738 pixels were finally available to populate the training and testing datasets (Table 5). The LITERAL dataset that represented only a small fraction of the available patches over wheat crops was kept for testing. The PHENOMOBILE dataset was split randomly between training (30%) and testing (70%) datasets (Table 5), resulting in 1803 pixels used to train the SVM model. Similarly, P2S2 was randomly split into 4329 pixels for training (about 40%) and the remaining for testing. This allows to get a balanced distribution between the contributions of PHENOMOBILE and P2S2 datasets to the training process, and Green/Senescent pixels fractions.

**Table 5:**
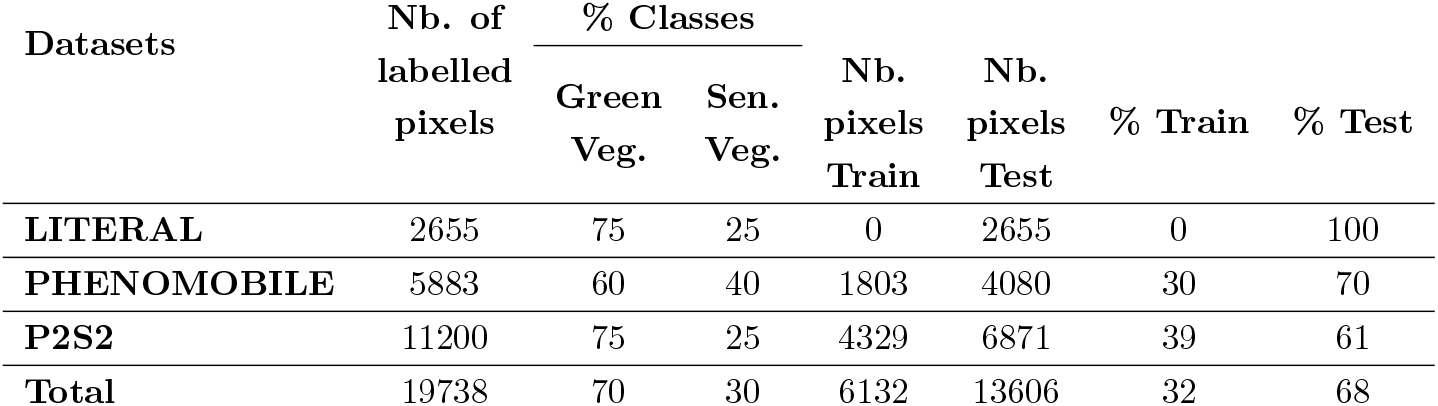
Distribution of the labeled pixels into the training and testing datasets. Only the pixels labeled as ***Green Veg***., ***Sen. Veg***., were used.

| Datasets | Nb. of<br>labelled<br>pixels | % Classes |  | Nb.<br>pixels<br>Train | Nb.<br>pixels<br>Test | % Train | % Test |
| --- | --- | --- | --- | --- | --- | --- | --- |
|  |  | Green<br>Veg. | Sen.<br>Veg. |  |  |  |  |
| **LITERAL** | 2655 | 75 | 25 | 0 | 2655 | 0 | 100 |
| **PHENOMOBILE** | 5883 | 60 | 40 | 1803 | 4080 | 30 | 70 |
| **P2S2** | 11200 | 75 | 25 | 4329 | 6871 | 39 | 61 |
| **Total** | 19738 | 70 | 30 | 6132 | 13606 | 32 | 68 |

### 2.4 Evaluation metrics

Since semantic segmentation classifies each individual pixel, three standard classification metrics derived from the confusion matrix were used to quantify the performances of the methods at the class level: precision, recall, and F1-score (Table 6). Further, the overall accuracy and overall F1-score were also computed to get a more global evaluation of the segmentation performances (Table 6). We also considered the fraction of pixels from a certain class in an image in a given viewing direction. This trait is widely used as a proxy of crop development [46] particularly for the green parts characteristic of the photosynthetically active elements [47]. Finally, for regression purposes RMSE and R^2^ were also considered. All these metrics were computed over the test dataset (Table 5), either directly on the test pixels of the image grids, for grid canopy fractions, directly on images grids from which the training pixels have been removed, or finally on the whole images.

**Table 6:** Metrics used to evaluate the performances of the models

| Metrics | Name | Definition |
| --- | --- | --- |
| **True positive** | $Tp_{class}$ | Number of pixels well predicted in the given class |
| **True negative** | $Tn_{class}$ | Number of pixels well predicted as not in the given class |
| **False positive** | $Fp_{class}$ | Number of pixels wrongly predicted in the given class (confusion) |
| **False negative** | $Fn_{class}$ | Number of pixels wrongly predicted as not in the given class (missing pixels) |
| **Precision** | $Prec_{class}$ | $\frac{Tp}{Tp + Fp}$ |
| **Recall** | $Rec_{class}$ | $\frac{Tp}{Tp + Fn}$ |
| **Accuracy** | $Acc_{class}$ | $\frac{Tp + Tn}{Tp + Tn + Fp + Fn}$ |
| **F1<sub>score</sub>** | $F1_{class}$ | $\frac{2 * Tp}{2 * Tp + Fp + Fn}$ |
| **Overall F1<sub>score</sub>** | $F1_{All}$ | $\frac{1}{N} \sum_{i=0}^N F1 - score_i$ |
| **RMSE** | RMSE | $\sqrt{(\frac{1}{n}) \sum_{i=1}^n (y_i^{theoretical} - y_i^{predicted})^2}$ |
| **R<sup>2</sup>** | $R^2$ | $1 - \frac{\sum (y_i^{predicted} - y_i^{theoretical})^2}{\sum (y_i^{theoretical} - y_i^{mean})^2}$ |
| **Canopy Fraction** | $CF_{class}$ | $\frac{\sum_{i=1}^I \sum_{j=1}^J (image(i, j) = class)}{\sum_{i=1}^I \sum_{j=1}^J (image(i, j))}$<br>(Where I and J are respectively the width and height of the image in pixels.) |

## 3 Results

### 3.1 Performances of the SegVeg model

#### 3.1.1 Separation of Vegetation | Background with the U-net first stage model

Results (Table 7) show that the U-net first stage model classifies well the vegetation from the background pixels, with an overall mean F1 score between 82% and 90%. The F1class values are higher for the Vegetation class. The confusing parts are mainly the distinction between the Vegetation on the ground and the ground itself (Fig. 5, bottom), or its equivalent confusion, i.e. senescence considered as ground residues (Fig. 5, top). The P2S2 sub-datasets, achieved the best F1all performances.

**Table 7:**
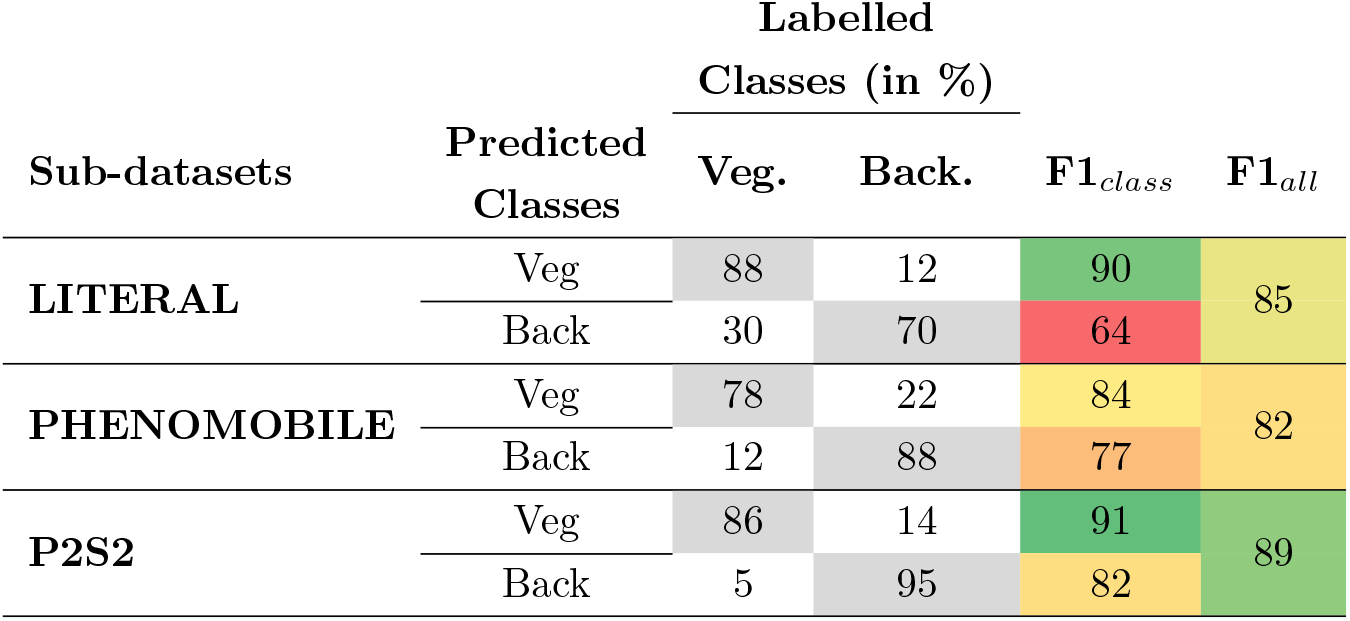
Performances of the U-net model used in SegVeg to classify vegetation (Veg.) and background (Back.) pixels. The elements of the confusion matrix, F1class and F1all are presented.

| Sub-datasets | Predicted<br>Classes | Labelled<br>Classes (in %) | | $F1_{class}$ | $F1_{all}$ |
| --- | --- | --- | --- | --- | --- |
|  |  | Veg. | Back. |  |  |
| **LITERAL** | Veg | 88 | 12 | 90 | 85 |
|  | Back | 30 | 70 | 64 |  |
| **PHENOMOBILE** | Veg | 78 | 22 | 84 | 82 |
|  | Back | 12 | 88 | 77 |  |
| **P2S2** | Veg | 86 | 14 | 91 | 89 |
|  | Back | 5 | 95 | 82 |  |

**Figure 5:**
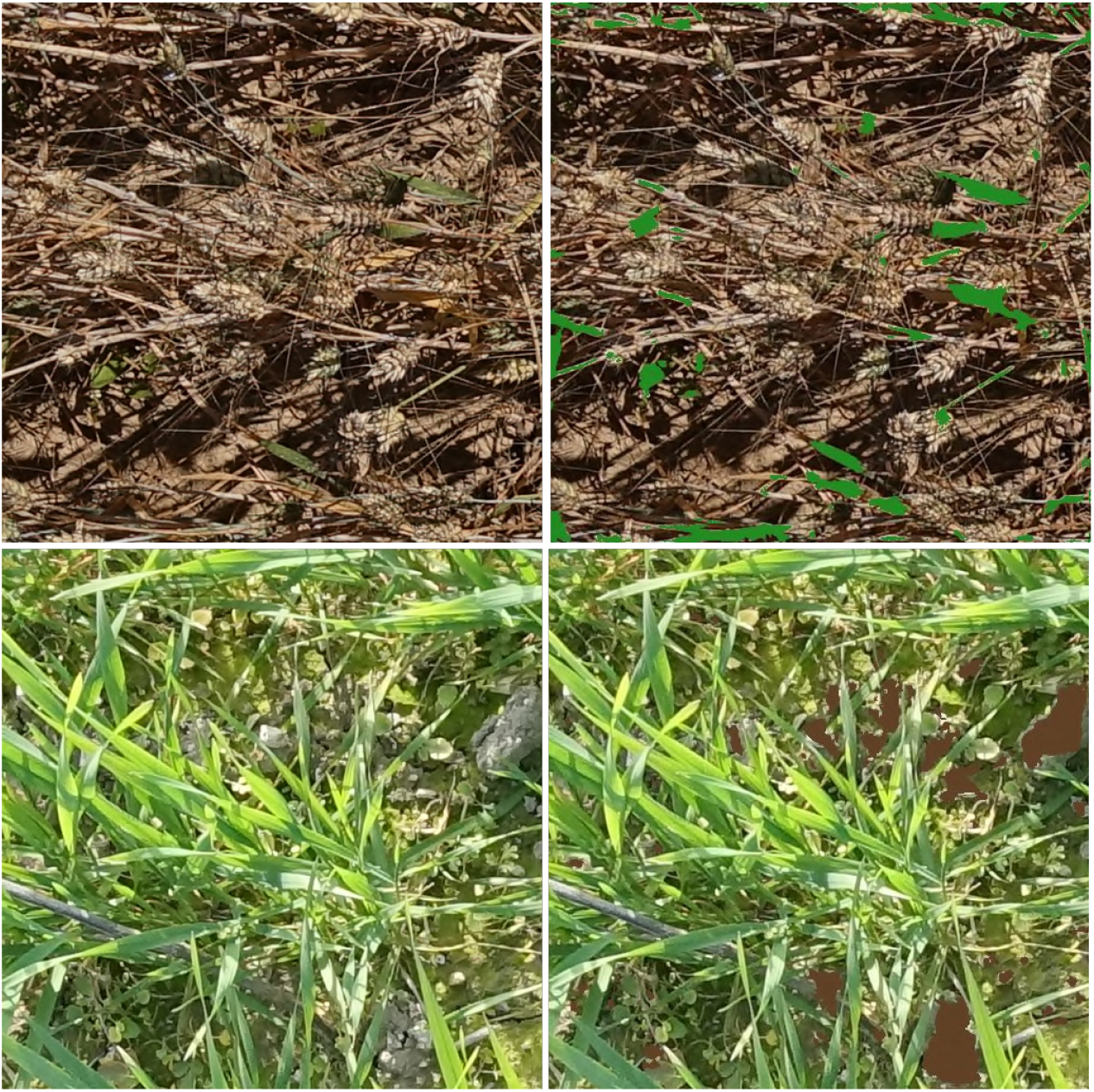
Examples of Vegetation | Background U-net first stage model confusions between ground and vegetation. On top, Vegetation considered as Background. On bottom, Soil considered as Vegetation. Green surfaces are pixels defined as Vegetation. Brown surfaces are pixels defined as Background. As model is binary everything else is considered as the opposite class.

#### 3.1.2 Classifying the green and senescent vegetation using a SVM

The SVM trained over the vegetation pixels was applied to all the pixels of the patches. Results (Table 8) show that the green vegetation pixels are generally well identified for the three sub-datasets.

**Table 8:** Confusion matrix (in % of the labelled pixels), Accuracy, and F1all values computed for the SVM classification alone and using the SegVeg model for the three sub-datasets. The diagonal terms of the confusion matrix are indicated in gray color. The colors of the two last columns correspond to the accuracy and F1all values (dark green, highest; dark red, lowest).

| Sub-dataset | Model | Predicted<br>Classes | Labelled Classes (%) |  |  | Acc | F1<sub>all</sub> |
| --- | --- | --- | --- | --- | --- | --- | --- |
|  |  |  | Sen. Veg. | Green Veg. | Back. |  |  |
| **LITERAL** | SVM alone | Sen. Veg. | 78 | 4 | 37 | 74 | 52 |
|  |  | Green Veg. | 22 | 96 | 63 |  |  |
|  | SegVeg | Sen. Veg. | 82 | 6 | 13 |  |  |
|  |  | Green Veg. | 9 | 93 | 26 | 84 | 79 |
|  |  | Backg. | 9 | 1 | 60 |  |  |
| **PHENOMOBILE** | SVM alone | Sen. Veg. | 90 | 4 | 61 | 60 | 47 |
|  |  | Green Veg. | 10 | 96 | 39 |  |  |
|  | SegVeg | Sen. Veg. | 61 | 3 | 5 |  |  |
|  |  | Green Veg. | 5 | 89 | 7 | 83 | 80 |
|  |  | Backg. | 33 | 8 | 87 |  |  |
| **P2S2** | SVM alone | Sen. Veg. | 89 | 2 | 54 | 69 | 48 |
|  |  | Green Veg. | 11 | 98 | 46 |  |  |
|  | SegVeg | Sen. Veg. | 77 | 2 | 6 |  |  |
|  |  | Green Veg. | 5 | 96 | 6 | 91 | 86 |
|  |  | Backg. | 18 | 2 | 88 |  |  |

The senescent vegetation pixels are more difficult to identify, with significant confusion with the green vegetation for the LITERAL sub-dataset. The background pixels are preferentially classified as senescent vegetation, except for the LITERAL sub-dataset (Table 8). This highlights the importance of the separation between the vegetation and the background since using only the RGB color information, i.e. without contextual information, appears not sufficient to separate well vegetation from background pixels, particularly for the senescent vegetation and the dark pixels as illustrated in Figure 6.

**Figure 6:**
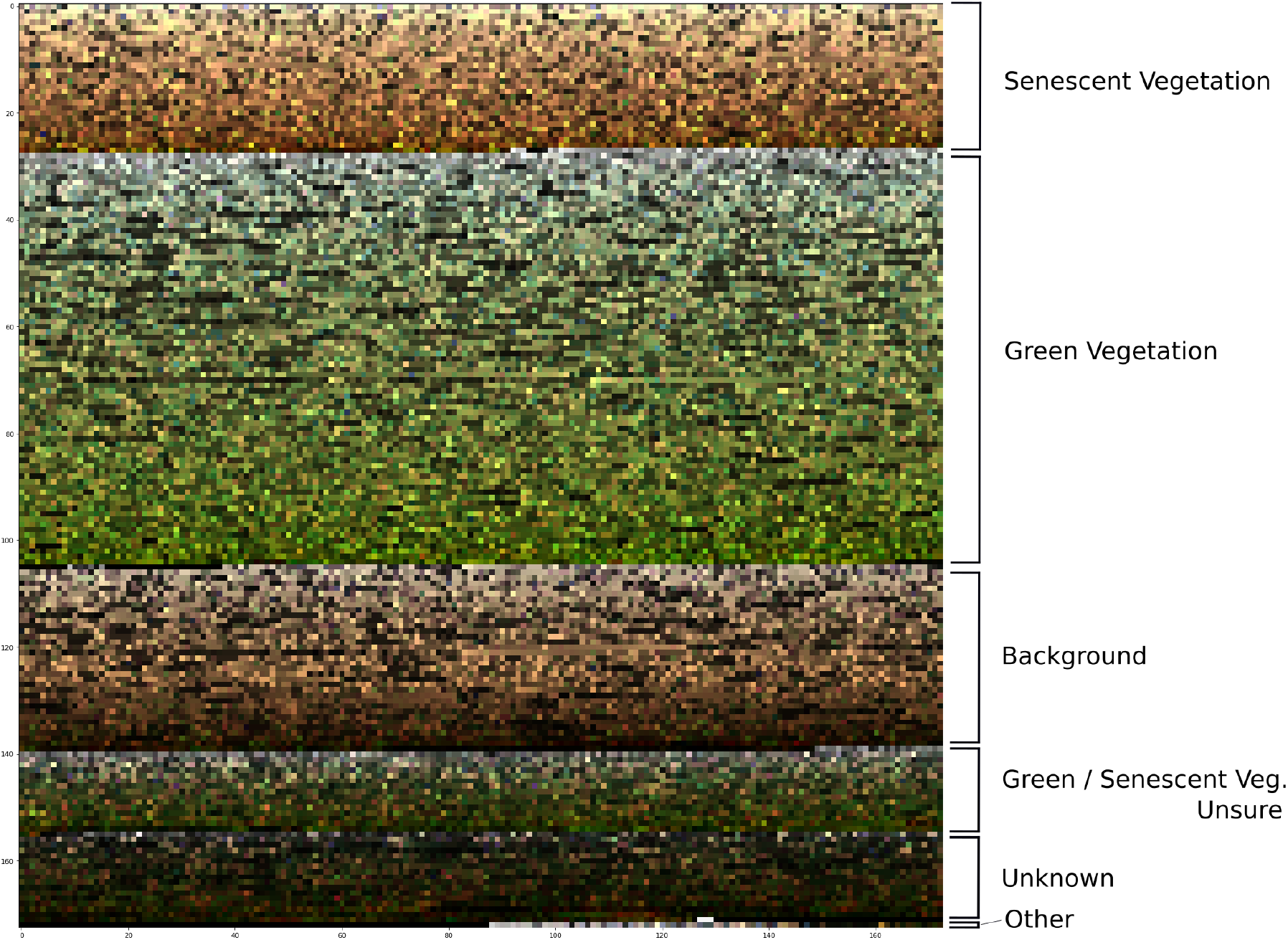
Distribution of the colors among the six classes as observed over the labeled pixels of the test and training datasets. For each class, pixels are sorted according to their brightness from the HSV color space

#### 3.1.3 Performances of the SegVeg model

The U-net Vegetation/Background segmentation followed the SVM classification of the vegetation pixels into green and senescent results in the SegVeg model. Results obtained on the test dataset show that the Accuracy and F1all score of the SegVeg model are high for the three sub-datasets. The SegVeg model classifies generally well the pixels into the three classes (Table 8, Fig. 7) because of the good performances of the two stages demonstrated earlier.

**Figure 7:**
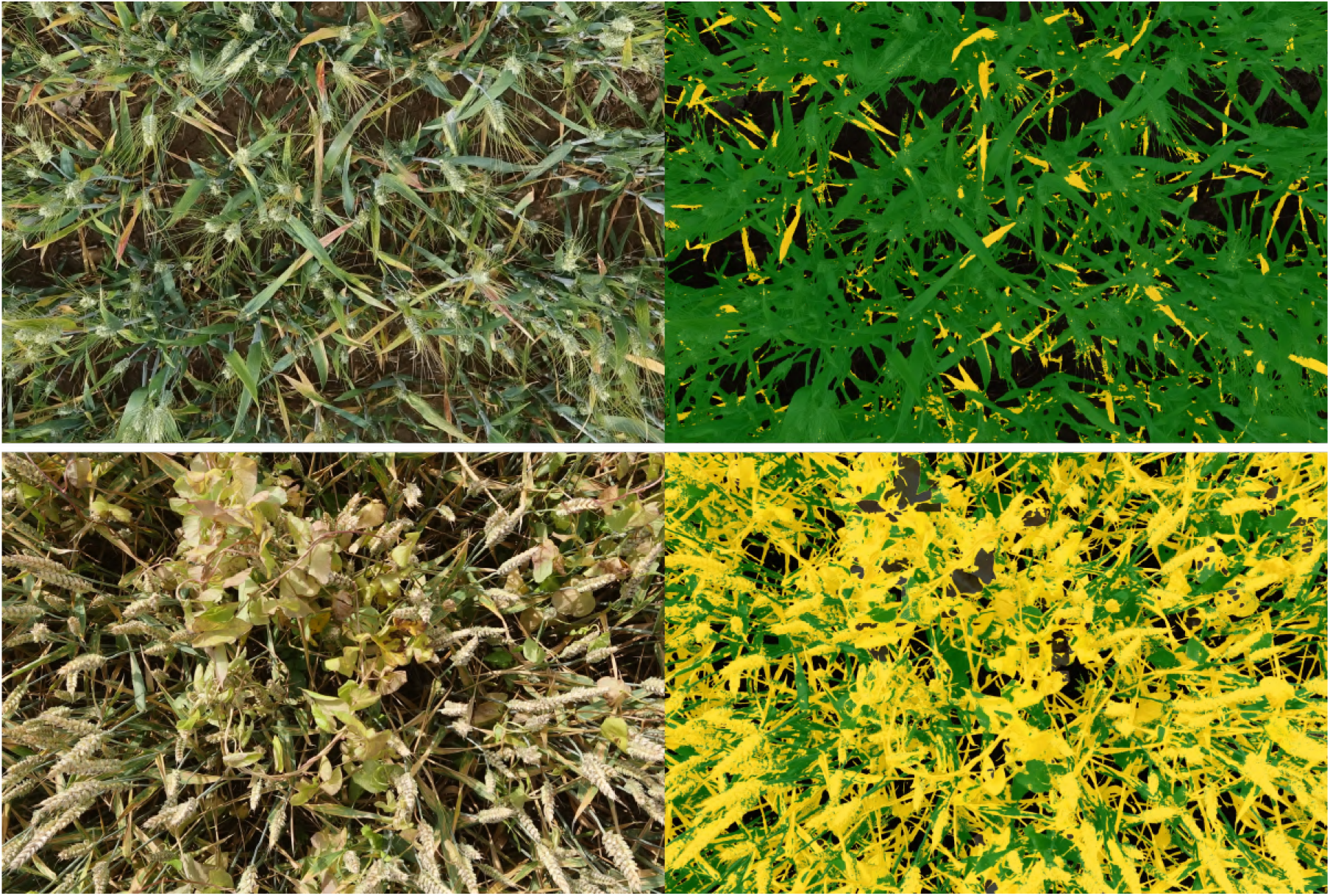
Examples of SegVeg model predictions on whole images wheat data from LITERAL system on early and late senescence stage. On the left, the original RGB images. On the right, the corresponding segmented images where the background green vegetation and senescent vegetation are represented respectively in black, green and yellow.

However, some significant confusion is still observed between the senescent vegetation and the background for the PHENOMOBILE dataset, and between the background and the green vegetation for the LITERAL dataset (Table 8). The degraded performances observed on the LITERAL dataset could be explained by the complexity of the images due to presence of awns that are smaller than the pixel size, inducing confusion between classes (Fig. 8).

**Figure 8:**
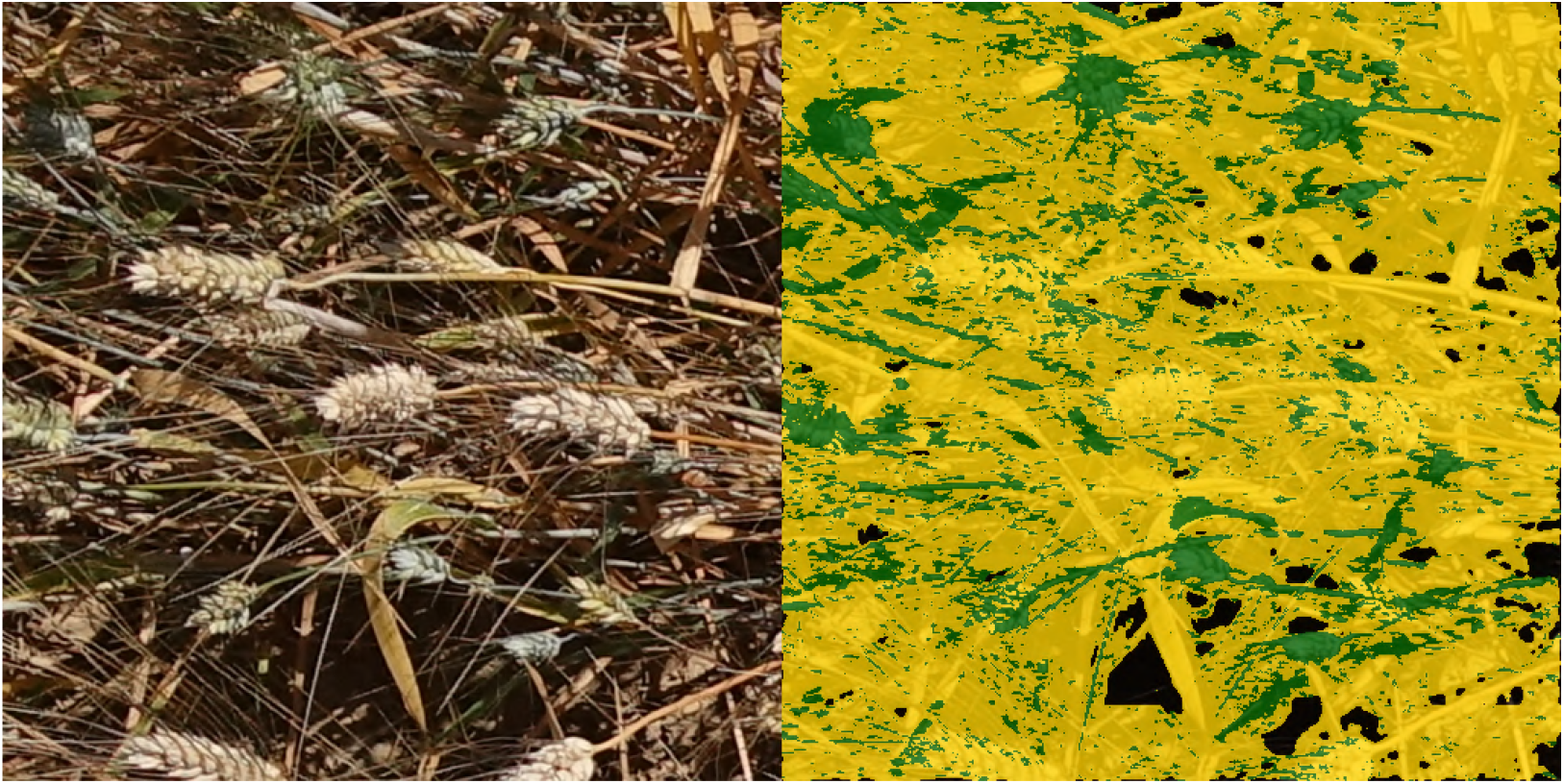
Examples of SegVeg model confusions on a complex LITERAL image with lots of thin spikes.

The classification performances of SegVeg seem to slightly degrade when the green fraction decreases and when the senescent fraction increases (Figure 9). These situations are underrepresented in the training database, which may contribute to the degraded performances observed.

**Figure 9:**
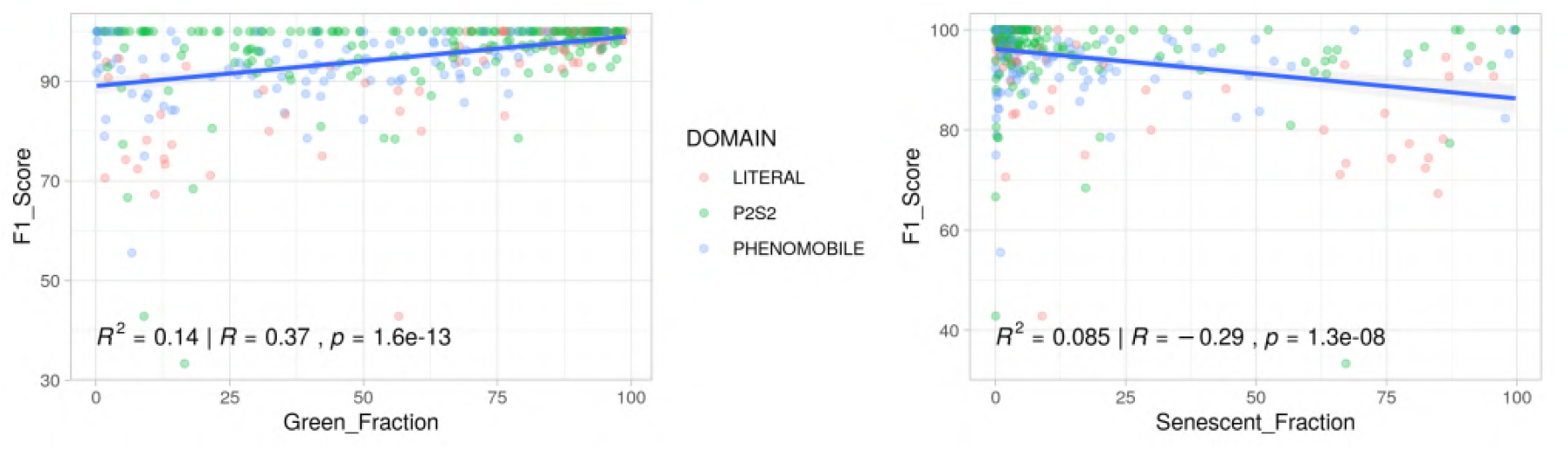
Performances (F1all) of SegVeg model as a function of the green and senescent fraction per image

### 3.2 Comparison with the U-net 3C

Results show that the U-net 3C (Table 9) performs closely to SegVeg (Table 8) on the three datasets pixels grids. The *Similitude* between the two models has been further studied by looking at the correspondence between the predicted classes (Table 9). The average Accuracy and F1all values for the similitude are respectively 90 and 85, with high values of the diagonal terms of the confusion matrix. However, on the average, the SegVeg model get slightly higher Accuracy (86) and F1all (84) values as compared to U-net 3C that gets respectively 82 and 79 for Accuracy and F1all. The confusion matrix (Table 8 and 9) reveals that best performances for SegVeg comes mostly from a better identification of the background pixels, particularly for the LITERAL dataset. The green vegetation is the best predicted by both models (Table 8 and 9). The senescent vegetation is slightly confounded with the background, while the background is about equally confounded with both vegetation classes (Table 8 and 9). For both models, the P2S2 sub-dataset achieves the best performances, and the LITERAL the worst ones, particularly die to higher confusion for the background class (Table 8 and 9).

**Table 9:** U-net 3 class and similitude to SegVeg model evaluation (in %). * for the Similitude, the labelled classes correspond to the classes predicted by the SegVeg model. The diagonal terms of the confusion matrix are indicated in gray cells. The color of Accuracy and F1all are related to their values (dark red the lowest; dark green the highest).

| Sub-dataset | Model | Predicted<br>Classes | Labelled Classes (%) * |  |  | Acc | F1<sub>all</sub> |
| --- | --- | --- | --- | --- | --- | --- | --- |
|  |  |  | Sen. Veg. | Green Veg. | Back. |  |  |
| LITERAL | U-net 3C | Sen. Veg. | 86 | 4 | 28 | 83 | 74 |
|  |  | Green Veg. | 9 | 95 | 31 |  |  |
|  |  | Backg. | 4 | 1 | 41 |  |  |
|  | Similitude | Sen. Veg. | 87 | 2 | 30 | 89 | 83 |
|  |  | Green Veg. | 12 | 67 | 10 |  |  |
|  |  | Backg. | 1 | 1 | 60 |  |  |
| PHENOMOBILE | U-net 3C | Sen. Veg. | 67 | 3 | 8 | 83 | 81 |
|  |  | Green Veg. | 12 | 89 | 7 |  |  |
|  |  | Backg. | 21 | 8 | 84 |  |  |
|  | Similitude | Sen. Veg. | 76 | 3 | 11 | 87 | 83 |
|  |  | Green Veg. | 13 | 94 | 5 |  |  |
|  |  | Backg. | 11 | 4 | 83 |  |  |
| P2S2 | U-net 3C | Sen. Veg. | 74 | 3 | 6 | 90 | 85 |
|  |  | Green Veg. | 11 | 95 | 6 |  |  |
|  |  | Backg. | 14 | 3 | 87 |  |  |
|  | Similitude | Sen. Veg. | 80 | 2 | 4 | 93 | 90 |
|  |  | Green Veg. | 15 | 96 | 3 |  |  |
|  |  | Backg. | 6 | 2 | 93 |  |  |

## 4 Discussion

### 4.1 Use color spaces to better describe the green vegetation

Differences of the sensitivity between operators impact the perception of color [48] and may therefore induce disagreement between them. Further, first stages of senescence may also create differences between the labeling of operators, since the yellow and reddish colors observed are in continuity with the green ones in the color space. To account for this effect, the labeling was done using several operators to get more consensus labeling.

The colors identified as senescent vegetation during the SVM classification of the vegetation pixels show that simple thresholds in the RGB space are not sufficient to get a good separation. Reciprocally, the same applies to the green vegetation. The combined use of some components of other color representation seem to be useful to segment the green vegetation as proposed by other authors such as [R, S, a, b, Cb, Cr] in [20], sRGB space used for CIELab transformation, in [49], or [H, S] in [50]. However, additional features may be also used to more specifically separate the senescent including the CMYK color space or the quadrature from YIQ that were selected as input features of the SVM because of their ability to indicate the senescent parts of vegetation (Figure 10).

**Figure 10:**
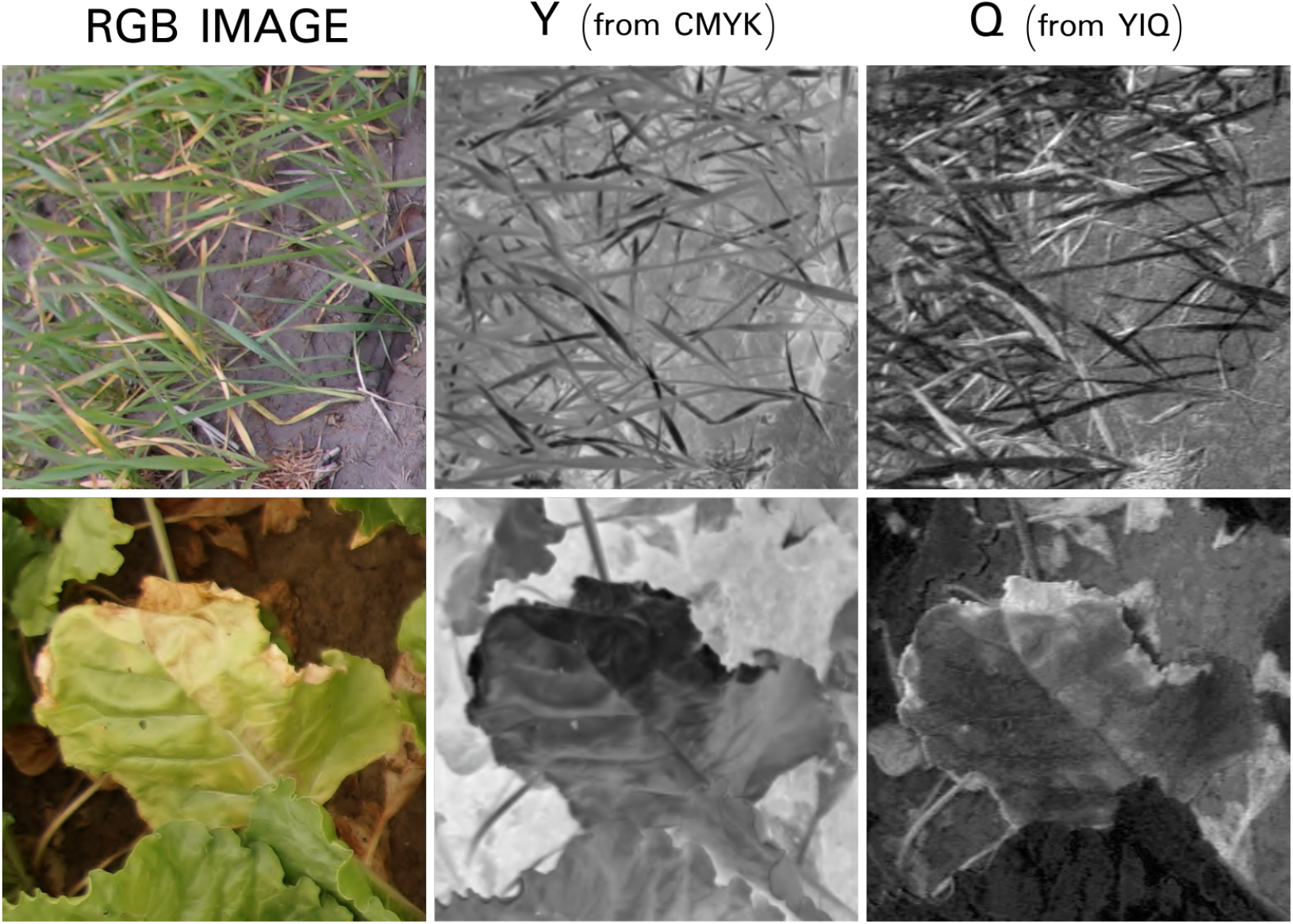
Output of different meaningful color spaces, respectively, RGB original image (left), Y component from CMYK, and Q component from YIQ (right). Y and Q images are in gray scale.

To review the model performances using these features and the corresponding boundaries, a 3D space RGB cube of 35^3^ voxels was created, containing then a huge panel of colors shades, which will help to discern where SegVeg model estimates senescent vegetation (Figure 11).

**Figure 11:**
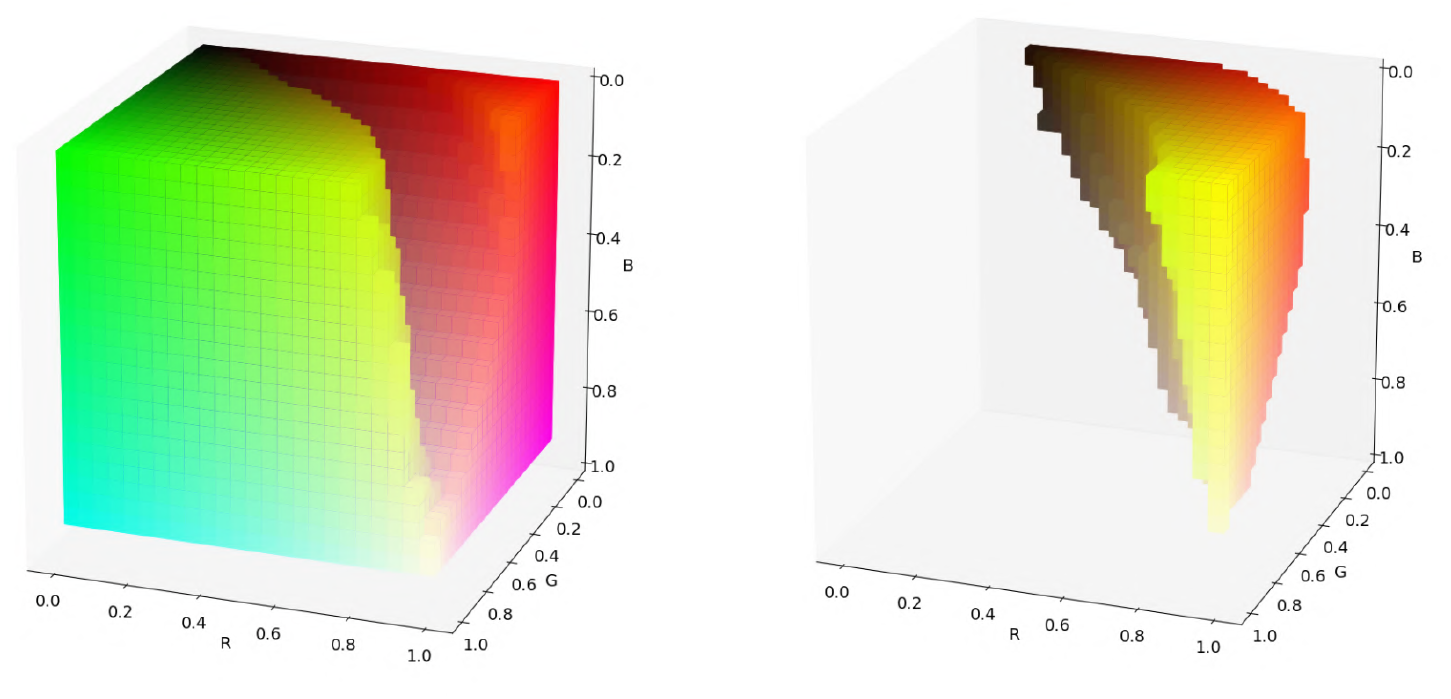
Boundaries of SegVeg colors inferred on a 35^3^ voxels RGB cube thanks to binary SVM of SegVeg second stage. On right, yellow predicted pixels. On left, the rest i.e including green predicted pixels.

### 4.2 Impact of illumination conditions on the segmentation performances

The misclassified pixels by the SegVeg model (Fig. 12, left) corresponds mostly to brownish colors representative of the senescent or background pixels. The few green pixels observed for high brightness and saturation may correspond either to errors in the labeling, or to pixels very close to the limit between the green and senescent vegetation (Fig. 11). Illumination conditions may strongly impact the quality of the classification. Misclassified pixels are preferentially observed for the small brightness values (Fig. 12, right) where the dynamics of the color values may be too limited to get an accurate classification based both on the color spaces or on the spatial features, inducing confusions between the three classes. This applies both to the labeling process and to the model predictions. Misclassified pixels are also observed preferentially for the highest brightness values (Fig. 12, right). In such conditions, some authors [51], propose to assign the saturated pixels to the most frequently saturated class. However, in our case, this would degrade the segmentation performances since the saturated pixels may belong to the three classes, with however a larger representation of green vegetation particularly with glossy leaves under either clear sky conditions or using flashes.

**Figure 12:**
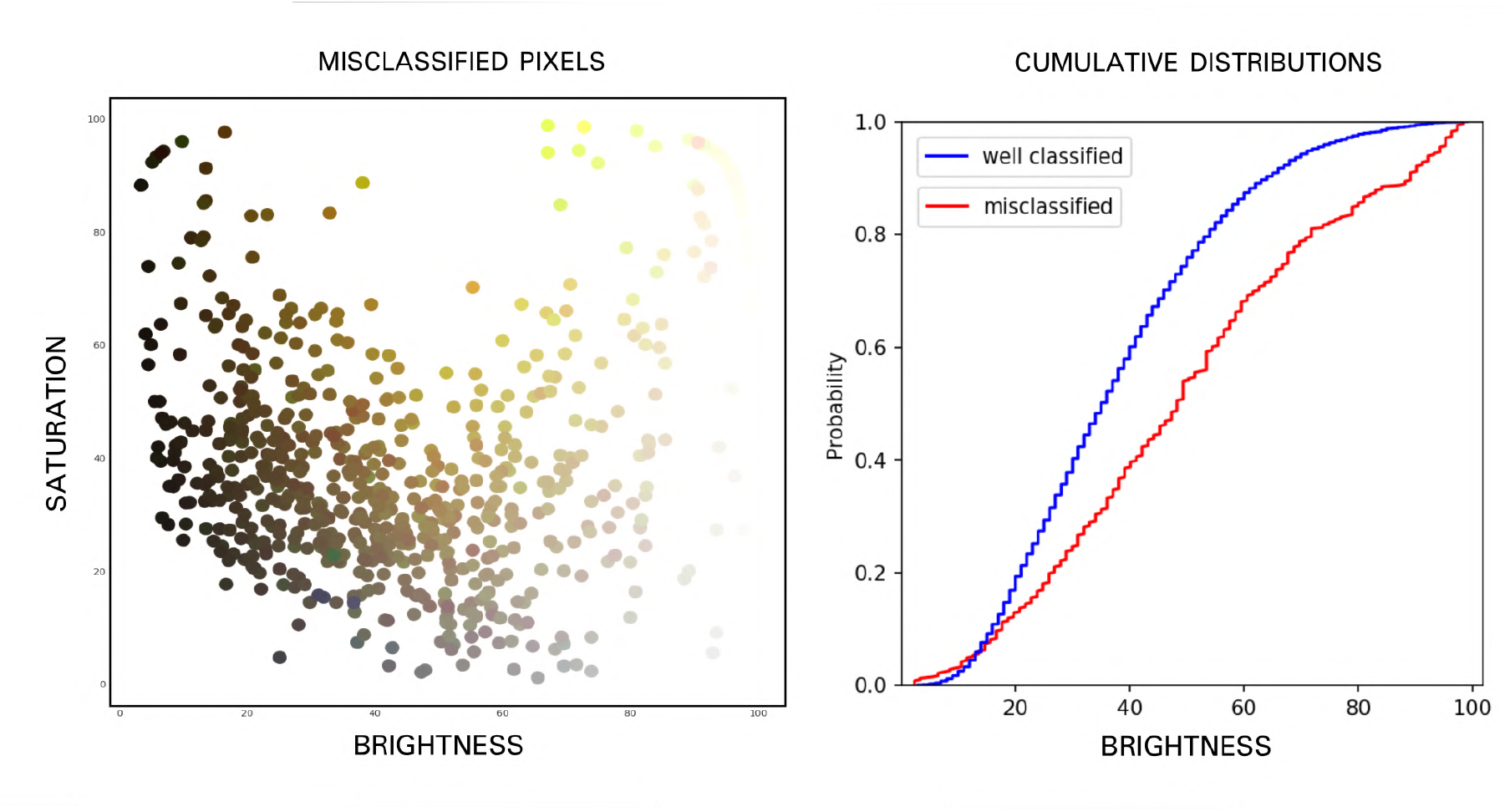
On the left, distribution of the Brightness (V from HSV) and Saturation (S from HSV) for the misclassified pixels by the SegVeg model. Each point corresponds to a mis-classified pixel from the grids of the test dataset. They are represented by their actual RGB color. Cumulated distribution of the brightness (right) of the misclassified (red) and well-classified (blue) pixel

The confusions observed for the PHENOMOBILE sub-dataset and leading to slightly degraded segmentation performances (Figure 13, bottom) are partly due to the use of flashes instead of the natural illumination for the LITERAL and P2S2 sub-datasets. The non-collimated nature of the light emitted by the flashes induces a decrease of the intensity of the radiation that varies as the inverse of the square of the distance to the source. When the source is too close to the top of canopy, pixels tend to be saturated with limited classification potentials. To limit this saturation effect, images taken from the PHENOMOBILE were slightly underexposed. Further, the pixels located at the bottom of the scene receive only very little illumination and are therefore very dark. The distribution of the brightness for the PHENOMOBILE (Figure 13, top) shows more darker pixels than the sub-datasets acquired under natural illumination conditions. This is in agreement with the larger confusion between the vegetation and the background presented earlier (Table 7,8).

**Figure 13:**
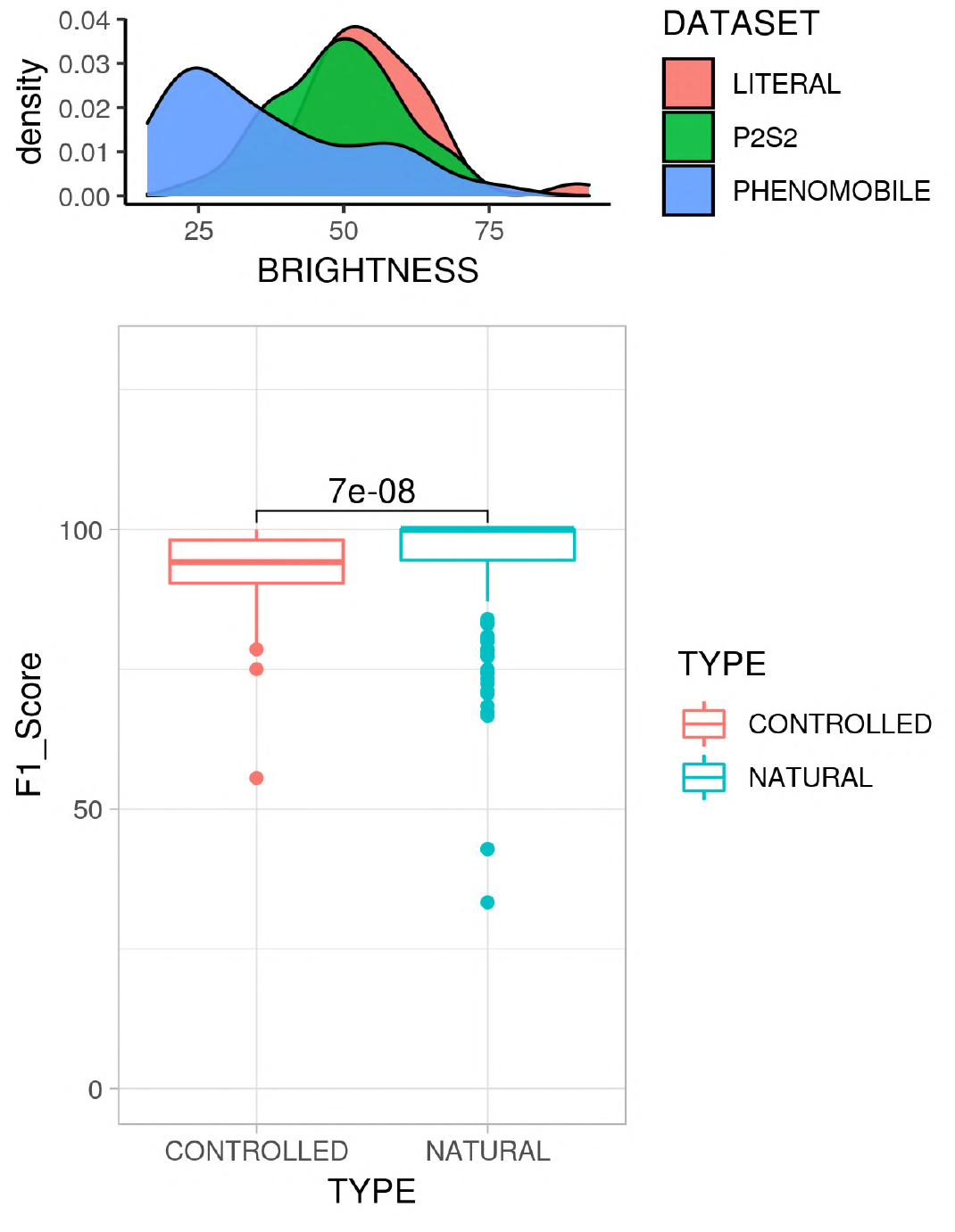
On the bottom, distribution of the performance (F1all) for both controlled (PHENOMOBILE) and natural (P2S2 and LITERAL datasets) illumination conditions. On top, distribution over brightness (V from HSV) of the three datasets.

### 4.3 Weak supervision is efficient

Because of the unavailability of images fully labeled into the three classes, U-net 3C was trained over masks predicted by SegVeg. This weak supervision approach could lead to biased predictions, since SegVeg predicted masks are not perfect as demonstrated previously in Table 8. Moreover, U-net 3C was trained over whole images compared to 6132 pixels for SVM classification model. However, the performances of U-net 3C (Table 9) are very close to those of SegVeg (Table 8) for the PHENOMOBILE and P2S2 sub-datasets, while SegVeg performs slightly better for the LITERAL sub-dataset. Comparison between SegVeg and U-net 3C (Table 9, “Similitude” case) confirms the good consistency between the two models. Weak supervision appears to be an efficient way to train Deep Learning algorithms by reducing the labelling process by the operators. The larger number of images therefore available to train the model is expected to partly compensate the lower quality of the “automatic” labelling. However, the main differences lie in the patterns of the green and senescent vegetation masks (Figure 14) where SegVeg appears crispier than U-net 3C that shows fuzzier masks. Indeed, the kernel filters used in U-net 3C to separate the green from the senescent vegetation tend to omit the small elements in the images and render diffuse more diffuse patches. Conversely, the pixel-based separation between the green and senescent vegetation allows to better describe the small details (Figure 14).

**Figure 14:**
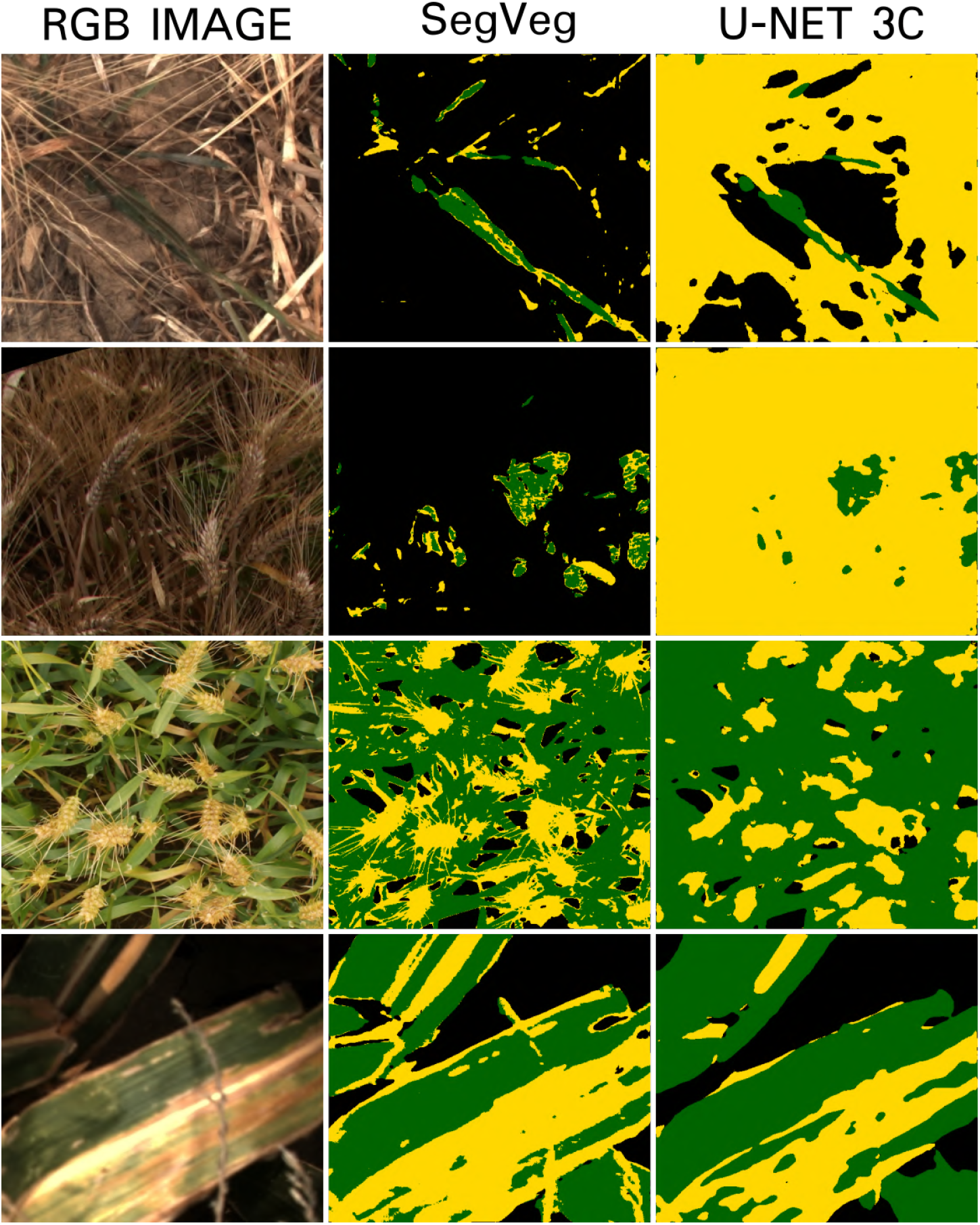
Results of the segmentation using SegVeg (middle) or U-net 3C (right). Bckground, Green vegetation and senescent vegetation are represented respectively in black, green and yellow. On the left, the original RGB image.

### 4.4 Predicting the fraction of green and senescent vegetation

The evaluation of the performances is completed over the grids that have been labelled by the operators. This corresponds first to a sub-sample of the image which questions the representativeness of the grid with regards to the entire image. We therefore evaluated the agreement between the segmentation predicted by SegVeg and that predicted by U-net 3C over both the grids and the images. Results show (Table 10, “Similitude” case) that the R^2^, RMSE, slope and offset for the grids and the images are in good agreement for each of the three fractions considered. This indicates that the fraction of background, green vegetation and senescent vegetation computed over the grid sub-sampling represents well the whole image.

**Table 10:** Performances of SegVeg and U-net 3C to estimate the background, Green vegetation and senescent vegetation fractions. The model “Similitude” corresponds to the comparison between SegVeg and U-net 3C. The performances were computed using either the labeled grids or on the whole images (U-net 3C testing images) for “Similitude”. R^2^ is the determination coefficient. The color of R^2^ and RMSE are related to their column values (dark green the best; dark red for the worst).

| Fraction | Model | Grid/Image | $R^2$ | RMSE | Slope | Offset (abs.) |
| --- | --- | --- | --- | --- | --- | --- |
| **Background** | SegVeg | Grid | 0.73 | 0.14 | 0.97 | 0.01 |
|  | U-net 3C | Grid | 0.74 | 0.14 | 0.99 | 0.01 |
|  | Similitude | Grid | 0.82 | 0.13 | 0.92 | 0.00 |
|  |  | Image | 0.80 | 0.13 | 0.94 | 0.01 |
| **Green Veg.** | SegVeg | Grid | 0.94 | 0.08 | 0.99 | 0.00 |
|  | U-net 3C | Grid | 0.90 | 0.10 | 1.03 | 0.01 |
|  | Similitude | Grid | 0.95 | 0.07 | 1.04 | 0.01 |
|  |  | Image | 0.95 | 0.07 | 1.07 | 0.02 |
| **Sen. Veg.** | SegVeg | Grid | 0.70 | 0.13 | 0.95 | 0.00 |
|  | U-net 3C | Grid | 0.70 | 0.13 | 1.14 | 0.02 |
|  | Similitude | Grid | 0.74 | 0.14 | 1.02 | 0.00 |
|  |  | Image | 0.73 | 0.13 | 1.07 | 0.00 |

SegVeg and U-net 3C show very similar performances. Best agreement is observed for the green vegetation fraction (Table 10), with a slight advantage for SegVeg, confirming the slightly better performances of the segmentation for this class (Table 8 and 9). The estimates are not biased (Table 10 and Figure 15, left). The estimation of the background and senescent vegetation fractions show degraded performances related to the degraded performances also observed previously on the segmentation of these two classes. The confusion between the background and the senescent vegetation pixels may be very large as highlighted by many outliers, with a quasi-exact compensation between these two fractions since the green vegetation fraction is well predicted (Figure 15, right). Small biases are observed for these fractions predicted by SegVeg and U-net 3C models, except for the senescent fraction of U-net 3C showing a significant bias (Table 10) mostly coming from the distribution of the outliers (Figure 15).

**Figure 15:**
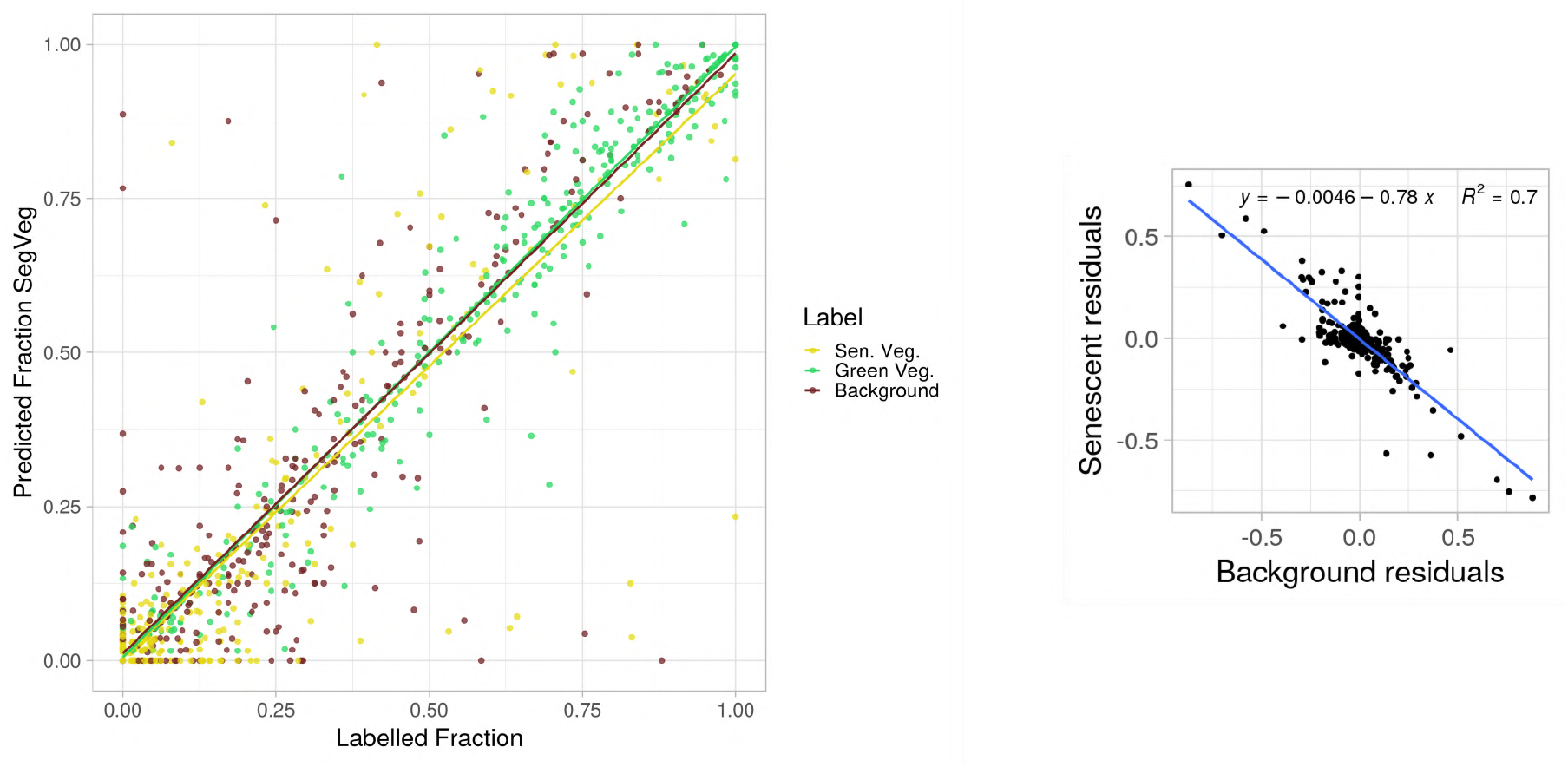
Comparison between the fractions predicted by the SegVeg model and the labelled ones. Each fraction is represented by a particular color. The best fit for each fraction is represented by a solid line of the same color.

The segmentation models SegVeg and U-net 3C appear efficient to compute the corresponding fractions. However, the SegVeg model offers a slight advantage regarding the better performances for the green fraction and smaller biases for the senescent vegetation fraction.

### 4.5 Limitation of the study

This study is based on segmentation models using shallow and deep learning techniques. It is therefore constrained by the available training and testing datasets. The first stage of the SegVeg model was using a relatively large and diverse training database (Table 1) containing 2015 images of 512 × 512 pixels. The second stage of SegVeg is trained over 6132 pixels extracted from grids applied to the original images also showing a wide diversity in species, stages, canopy state and acquisition conditions. However, the pixels labelled as uncertain (Green/Sen. Veg. Unsure, Unknown; Other) were not used, forcing the SVM model to extrapolate for these situations. The uncertain pixels represented about 16% of the labelled pixels (Table 4), with some variation across the three subdatasets. Further, the training was completed over two sub-datasets with the P2S2 overrepresented as compared to the PHENOMOBILE one. This is the reason why we presented the results per sub-dataset. Further, this explains also partly the differences of the performances observed between the three test datasets, with generally P2S2 *>* PHENOMOBILE *>* LITERAL.

The evaluation of the models was completed at the pixel level over the grids. A large number of pixels was considered here (13606 pixels, Table 5), including those extracted from the LITERAL subdataset that was not used for training the models. The “Unsure” pixels were not used to compute the performances, which may also induce small biases in the results since the “Unsure” pixels may not be evenly distributed between the three classes of interest. However, we did not have other alternatives. Since “Unsure” pixels correspond mostly to very dark, very bright (Figure 6) or to mixed pixels, great attention should be paid to the actual spatial resolution and the exposition of the images. Further, studies based on 3D scenes rendered realistically should be conducted to better understand the unsure classes and their possible distribution among the three classes of interest.

## Acknowledgments

This work received support from ANRT for the CIFRE grant of Mario Serouart, co-funded by Arvalis. The study was partly supported by several projects including ANR PHENOME (Programme d’investissement d’avenir ANR-11-INBS-0012)Digitag (PIA Institut Convergences Agriculture Numérique ANR-16-CONV-0004), CASDAR LITERAL and P2S2 funded by CNES. Many thanks to the people who annotated the datasets, including Frederic Venault and Micheline Debroux.

## Author Contributions

Mario Serouart and Simon Madec have written the code, analyzed the results and conducted reviews. Mario Serouart, Simon Madec, Kaaviya Velumani and Etienne David have conducted the pipeline. Frederic Baret, Marie Weiss and Raul Lopez Lozano have supervised experiments at all stages. All authors contributed to editing, reviewing, and refining the manuscript.

## Funding

The authors declare that there is no conflict of interest regarding the publication of this article.

## Data Availability

Upon acceptance of the paper, SegVeg pixels dataset, images and their corresponding segmentation masks will be publicly available. All the SegVeg scripts for computation and analysis are also public: https://github.com/mserouar/SegVeg. For simplicity, dataset download links (including Zenodo) will be specified in the above repository.

## Notes

https://github.com/mserouar/SegVeg

### Competing Interest Statement

The authors have declared no competing interest.

